# Optimizing crop varietal mixtures for viral disease management: A case study on cassava virus epidemics

**DOI:** 10.1101/2025.02.03.636186

**Authors:** C. Israël Tankam, Ruairí Donnelly, Christopher A. Gilligan

## Abstract

Cassava viral diseases, including Cassava Mosaic Disease (CMD) and Cassava Brown Streak Disease (CBSD), pose significant threats to global food security, particularly in sub-Saharan Africa. This study explores the potential of varietal mixtures as a sustainable disease management strategy by introducing CropMix, a novel web-based application. The application encodes a flexible insect-borne plant pathogen transmission model to predict and optimize yields under scenarios of varietal mixtures. For instance, we use the application to evaluate the ability of virus-resistant cassava varieties to protect more susceptible varieties against CMD and CBSD, and we also consider mixtures involving tolerant varieties and non-host crops. For CMD, the high transmission rates of cassava mosaic begomoviruses limits the efficacy of mixtures, with susceptible monocultures emerging as better than susceptible-resistant mixtures whatever the whitefly pressure. In contrast, for CBSD, varietal mixtures demonstrate substantial benefits, with resistant varieties shielding susceptible ones and mitigating severe yield losses under moderate or high insect pressure. Management strategies involving non-host crops and complementary control measures, such as roguing, can further enhance outcomes. The model’s simplicity and adaptability make it suitable for tailoring recommendations to diverse insect-borne crop viral diseases and agroecological contexts. The study emphasizes the need for integrating real-world data and participatory frameworks to refine and implement disease management strategies. We discuss the critical balance between agronomic potential and farmer acceptability, underscoring the importance of collaborative efforts to ensure sustainable cassava production.

## 1 Introduction

The prevalence of insect-borne diseases poses a significant threat to crop production worldwide, leading to substantial yield losses and economic hardship for farmers. In recent years, the emergence and spread of these diseases have escalated, fuelled by factors such as globalization, climate change, and the evolution of virulent pathogen strains [29]. Insect-borne viruses can infect plants, leading to a range of symptoms from mild leaf discolouration to severe stunting and even death. The use of genetically resistant plant varieties is fundamental to sustainable and effective control of plant pathogens[34]. The method is especially crucial for managing viral diseases, where resistance genes can disrupt viral replication, movement, or infection processes, offering long-term control [19]. Building on the characteristics of resistant varieties, cultivar mixtures can capitalize on differing protective traits to mitigate disease risks for susceptible crops.

Cultivar mixtures with varying resistance levels can improve the resilience of crops exposed to plant pathogens by interfering with pathogen spread, where resistant plants protect susceptible ones by reducing inoculum build-up. The approach reduces the risks of crop failure and maintains genetic diversity, essential for agricultural sustainability. For instance, studies suggest that effective mixtures can protect cereals against foliar diseases [14, 7, 6, 16]. This protective effect results from two main mechanisms: the barrier effect, which reduces transmission by reducing connectivity among susceptible plants, and the dilution effect, where resistant plants lower inoculum density. Cultivar mixtures not only help manage diseases but are also known to improve yield in cereal crops [2, 35]. Further study is needed to optimize their use across diverse crops and environments, including root, tuber, and banana (RTB) systems.

Among root and tuber crops (RTBs), cassava is highly susceptible to insect-borne viruses that cause Cassava Mosaic Disease (CMD) and Cassava Brown Streak Disease (CBSD). Both viruses are transmitted by the whitefly *Bemisia tabaci* and severely impact production across Africa. Cassava mosaic disease, caused by begomoviruses, results in leaf mosaics, stunted growth, and reduced yields [31, 25]. Cassava brown streak disease, caused by ipomovirus species, affects leaves and roots, leading to brown necrotic lesions in roots, which lowers yield and storage quality [17]. These diseases threaten cassava production and food security, particularly where cassava is a staple crop. Control methods include plant resistance, insecticides, agricultural practices include roguing, and clean seed (i.e. virus-free) systems [24, 33]. However, CBSD’s root impact complicates diagnosis, and CMD’s rapid spread challenges phytosanitation efforts [24], underscoring the importance of resistant cultivars. Resistance adoption and dissemination, notably for CMD, has progressed with CMD-resistant varieties from the International Institute of Tropical Agriculture (IITA) [23]. Cultivar mixtures can further aid disease control and yield, though selecting optimal proportions requires balancing complex variety interactions with environmental and pathogen factors. In this context, mathematical modelling offers a powerful tool for understanding the dynamics of disease transmission and optimizing mixture strategies.

Recent advances highlight mathematical and statistical modelling as valuable tools for understanding cultivar mixtures in cropping systems. Mikaberidze et al. [26] developed a susceptible-infected (SI) model of 2n equations to assess how host mixtures impact disease dynamics, showing that optimal mixtures reduce disease severity compared with pure stands, especially under partial pathogen specialization. Djidjou-Demasse et al. [8] extended the analysis, by modelling multi-locus gene-for-gene dynamics at both within- and between-cultivar scales. They showed that mixtures in the form of crop mosaics were generally more effective in reducing disease compared with pyramiding resistance genes within a single cultivar. The effect was most pronounced under high inoculum pressures. Clin et al. [4] examined pathogen virulence in relation to host diversity, establishing a minimum number of host varieties as crucial for effective disease management. Building on this, Clin et al. [5] analysed monovirulent and doubly virulent pathogen dynamics, suggesting optimal host ratios of respectively 30:70 or 70:30 and highlighting the roles of immune priming and resistance-breaking costs. Lastly, Hamelin et al. [15] explored the coexistence between wild-type and resistance-breaking pathogens, indicating that while host mixtures complicate disease dynamics, they offer sustainable management strategies. Together, these studies emphasize the potential of modelling to inform cultivar mixtures for improved disease control and crop resilience.

Despite these advances, gaps remain in understanding the role of vector behaviour in cultivar mixture dynamics, particularly in relation to disease spread. Modelling vector-borne diseases, especially insect-borne diseases, often overlooks the impact of vector behaviour on disease dynamics, focusing mainly on vector density rather than host-vector contact rates that are critical for viral transmission [32]. This oversight limits understanding of epidemic emergence and spread, prediction and control, particularly in cultivar mixtures. Here, we consider scenarios in which the rates of virus acquisition by virus-free vectors and the rates of transmission by viruliferous vectors vary by cultivar. We examine how the effects on inoculation and vector movement between varieties affect disease dynamics. We introduce a framework for modelling vector-borne plant viruses in a two-crop RTB mixture and present a software- informed approach to optimize variety mixtures for managing cassava mosaic disease in cassava crops. We prioritize yield optimization as small-scale farmers often focus on maximizing short-term economic returns from limited acreage rather than minimizing disease incidence. Yield improvement is immediate and measurable within a single season, aligning with farmers’ economic needs and fostering informed adoption of management options. Secondary consideration is given to the impacts on disease incidence. Along with resistance, we also analyse how tolerant varieties and decoy crops protect susceptible varieties. Tolerant varieties experience low yield loss despite severe infection, and decoy crops, though they don’t produce cassava tubers, provide a protective barrier against disease. The software developed for this study is freely available and built on the mathematical modelling framework.

## 2 Materials and methods

This section presents the mathematical framework for modelling disease dynamics, impacts on yield along with optimal variety mixture in response to insect-borne viral crop pathogens, with a focus on cassava viruses. The model integrates the dynamics of vector-borne transmission, within-host disease progression, plant variety mixtures, and roguing (removal of infected plants) as a management strategy. The goal is to optimize the mixture of cassava varieties to maximize crop yield. The framework is implemented in an application designed to provide decision support for farmers, researchers, and agricultural stakeholders, enabling real-time scenario analysis and output visualization.

### 2.1 Modelling: Assumptions and Process Overview

The mathematical model used to represent virus spread in a cassava field with two varieties (A and B) incorporates biological, epidemiological, and management-related assumptions. It simulates interactions between cassava plants and the whitefly vectors transmitting the virus, disease progression within plants, and management interventions such as roguing. The model describes field dynamics over a cropping season, focusing on two varieties with different virus susceptibilities and transmission rates. Figure 1 illustrates these processes.

**Figure 1:**
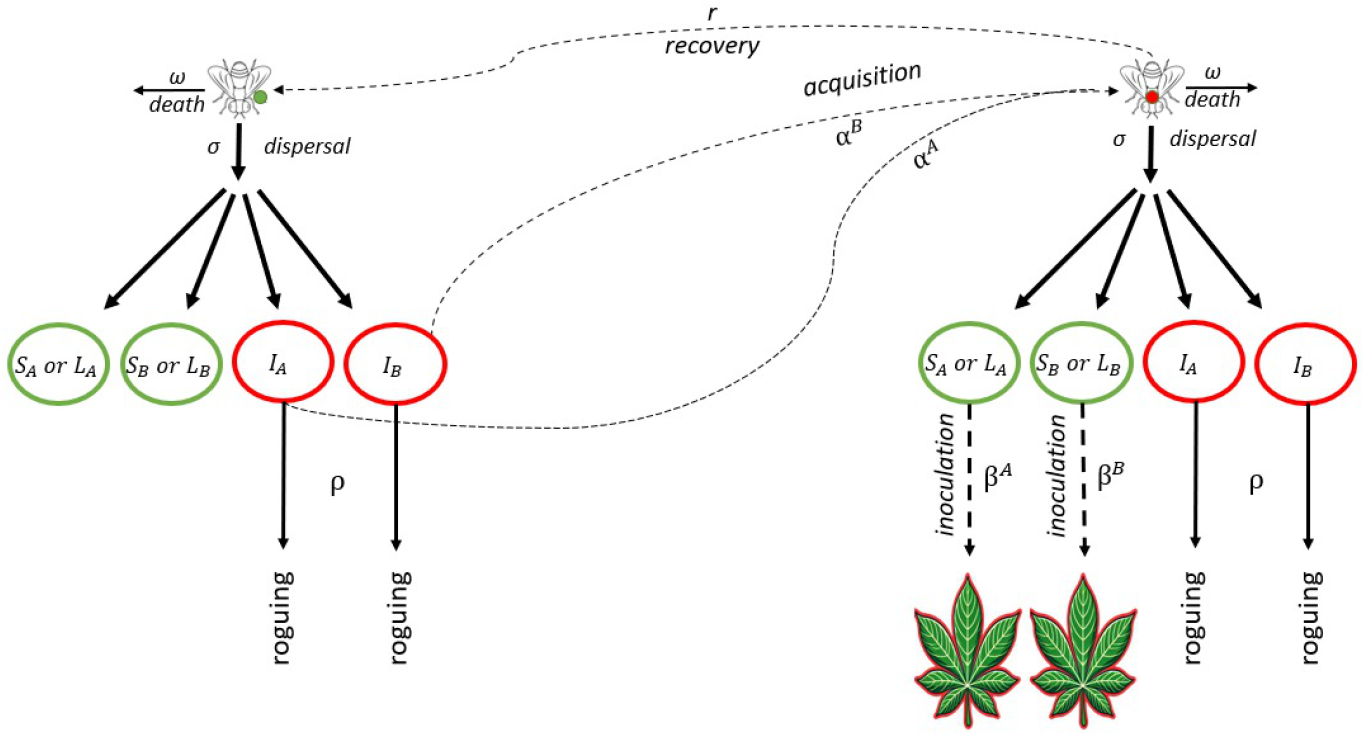
Schematic representation of insect vector behaviour in the context of varietal mixing and vector-borne pathogen transmission. The field consists of two types of plants, A and B, each having three possible disease states: (1) Healthy (*S*_*A*_ and *S*_*B*_), Latently infected (*L*_*A*_ and *L*_*B*_), and Infectious (*I*_*A*_ and *I*_*B*_). Upon dispersal, insect vectors settle on one of the six types of plants in proportion to their frequency. Virus-free vectors can acquire the virus from infectious plants at a rate that depends on the plant variety, while infectious vectors ultimately disperse the virus when they settle on a healthy plant, with an inoculation rate that also depends on the plant variety. Insect vectors experience the same dispersal and mortality rates regardless of virus-status. The schematic allows for regular roguing of infected plants, i.e., one of the principal means by which virus epidemics in host plants is managed.

As shown in the schematic (Figure 1), the field is planted with varieties A and B in proportions *θ* and 1−*θ*, respectively, maintaining a constant plant density *K*. Removed plants are replaced by susceptible plants of the same variety. Varieties differ in virus susceptibility and vector-related transmission rates. Non-viruliferous vectors can acquire the virus or die, being replaced by new vectors, while viruliferous vectors can die or recover, also replaced by non-viruliferous vectors. Once infected, plants progress through latent and infectious stages. Roguing removes infected plants at a rate *ρ*, with detection probabilities *d*_*A*_ and *d*_*B*_ determining removal rates. In this study, perfect detection is assumed (*d*_*A*_ = *d*_*B*_ = 1), ensuring immediate replacement of removed plants, and keeping the total density constant despite fluctuations in plant states.

The vector dynamics account for mortality (*ω*) and recovery (*r*), with recovered vectors returning to the susceptible compartment. Vectors move randomly among plants, and epidemic progression is tracked through healthy, latent, and infectious compartments. Over a fixed cropping season (*T* days), disease dynamics evolve, with infection, disease progression, and plant removal shaping outcomes. Final disease prevalence and yield are calculated based on remaining healthy and infected plants. Table 1 summarizes these processes, and Supplementary Material S1 provides detailed equations. This framework captures key dynamics of insect-borne disease spread in mixed-variety cassava fields and facilitates exploring management strategies. We now briefly describe the role of insect behaviour in pathogen transmission.

**Table 1:** Dynamics of virus transmission and vector interactions among plant compartments. The table summarizes the events involved in virus spread and vector behaviour, detailing the flow of individuals between compartments, along with the corresponding rates of each event. Each event is categorized by its type, including inoculation, incubation and roguing for plants of variety A or B; Acquisition, recovery, and insect mortality for infectious insect vectors that acquired the virus on either *A* plants (*V*^*A*^) or *B* plants (*V*^*B*^). The table summarises the interactions between healthy, latently infected, and infectious plant states, as well as the role of insect vectors in the transmission process. The inoculation rates *β*^*A*^ and *β*^*B*^ are in plant per vector per day, and the acquisition rates *α*^*A*^ and *α*^*B*^ are in vectors per day. All other event rates in the table are expressed per day. The overall insect abundance, F, in the field, assumed constant.

| Event | Flow between compartments | rate |
| --- | --- | --- |
| Inoculation | $S_A \longrightarrow L_A$ | $\beta^A \times \text{\#infectious vectors feeding on } S^A$ |
| | $S_B \longrightarrow L_B$ | $\beta^B \times \text{\#infectious vectors feeding on } S^B$ |
| Incubation | $L_A \longrightarrow I_A$ | $\gamma_A \times L_A$ |
| | $L_B \longrightarrow I_B$ | $\gamma_B \times L_B$ |
| Roguing | $I_A \longrightarrow S_A$ | $\rho \times \text{detection probability } d_A \times I_A$ |
| | $I_B \longrightarrow S_B$ | $\rho \times \text{detection probability } d_B \times I_B$ |
| Aquisition | $F - (V^A + V^B) \longrightarrow V^A$ | $\alpha^A \times \text{proportion of virus-free insects on } I_A$ |
| | $F - (V^A + V^B) \longrightarrow V^B$ | $\alpha^B \times \text{proportion of virus-free insects on } I_B$ |
| Recovery | $V^A \longrightarrow F - (V^A + V^B)$ | $r \times V^A$ |
| | $V^B \longrightarrow F - (V^A + V^B)$ | $r \times V^B$ |
| Insect mortality | $V^A \longrightarrow F - (V^A + V^B)$ | $\omega \times V^A$ |
| | $V^B \longrightarrow F - (V^A + V^B)$ | $\omega \times V^B$ |

### 2.2 Yield

Box1 summarizes the details on yield calculation and optimization. The goal is to determine the optimal proportion *θ* of variety A (and hence the optimal proportion 1−*θ* of variety B) in the field that results in the highest possible yield. For practicality and statistical significance, and to mitigate individual variability and sampling bias, yield measurements are conducted at the hectare level rather than at the individual plant level. We scale the model to a hectare. The field density *K* represents the planting density per hectare, typically ranging from 10,000 to 16,000 plants per hectare in cassava cultivation.

#### Box 1: Yield Calculation and Optimization

**Figure 2:**
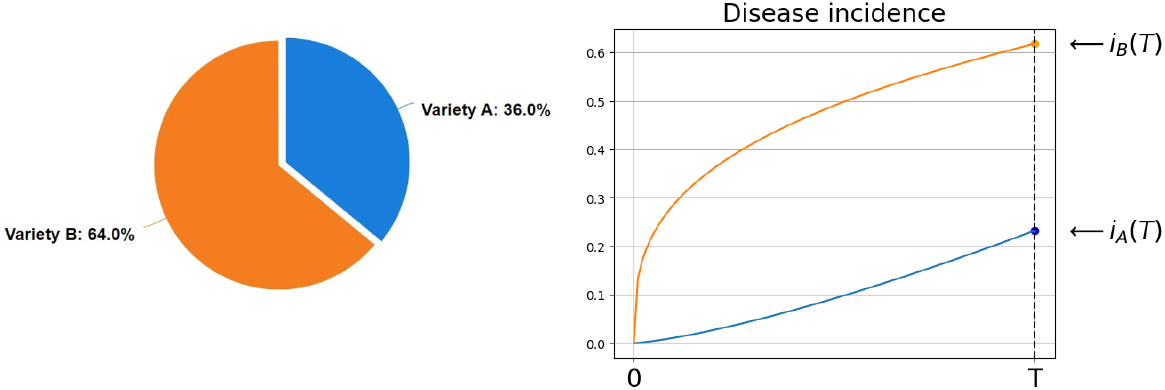
Illustration of a mixture and disease incidence dynamics for yield calculation

At the end of the growing season, a certain proportion of plants from each variety is infected, while the rest remain healthy. For variety A, which makes up a proportion of the total crop, the infected plants contribute to the yield according to how they perform under infection, while the healthy plants contribute according to their uninfected yield. Similarly, for variety B, the infected and uninfected plants contribute their respective yields based on their condition. The total yield of the mixed crop combines these contributions from both healthy and infected plants of each variety, considering the proportions of the crop and the infection levels at the end of the season. The total yield therefore reads:

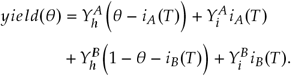

Where 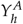 and 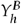 designate the yield when uninfected of variety A and B respectively, and (*θ* − *i*_*A*_(*T*))and (1 − *θ* − *i*_*B*_(*T*)) are their respective proportions.

**Optimization**

We aim to find the value of *θ* (the proportion of variety A in the mixture) that gives the highest total yield. This involves solving a problem where we maximize the yield, which is a smooth function of *θ* between 0 and 1. We do this as follows:

1. Start by testing values of *θ* from 0 to 1 in small steps, e.g. increments of 0.05.
2. For each tested value of *θ*, simulate how infection spreads through the crops over the season using the given model.
3. Calculate the total yield at the end of the season for each tested value of *θ*, considering the effects of infection and healthy plants’ contributions.
4. Find the *θ* value that produces the highest yield.
5. Once the best *θ* is identified, narrow the search to a smaller range around this value and test with smaller steps (e.g., 0.005) to refine the result and ensure accuracy.

This step-by-step process identifies the best proportion of variety A and B for maximizing the yield.

### 2.3 Cultivar Phytotypes

Thus far we have described a model of insect vector behavious incorporating pathogen transmission and we have indicated how yield may be optimised for a mixture of distinct host plant varieties. We now discuss the characterisation of host plant varieties. For simplicity, we refer to the main categories of varieties as phytotypes, including susceptible (*SUSC*), resistant (*RES*), tolerant (*TOL*) and decoy (*DEC*) phytotypes. Individual varieties are defined by the inoculation rate *α* (infected vector feeding on healthy plant), acquisition rate *β* (uninfected vector feeding on infected plant), yield when healthy (*Y*_*h*_), and yield when infected (*Y*_*i*_). For simplicity, for the purposes of illustrating the application with respect to management of cassava viruses, we assume equal rates of disease onset across plant varieties and equal detection probabilities for infected plants.

Our phytotypes include a susceptible (*SUSC*), resistant (*RES*), tolerant (*TOL*) and decoy (*DEC*). *SUSC*, the baseline, has parameter estimates for *α* and *β* that are based upon analysis of published laboratory data [9]. For the purposes of illustration, we characterise a typical resistant variety, as being defined by reduced acquisition and inoculation rates such that the value of *αβ* for *RES* is lower than that of *SUSC* by a factor of 90%, and in addition lower *RES* values for *Y*_*h*_ and *Y*_*i*_ than those for *SUSC* can be assumed reflecting a cost of resistance in terms of yield (note that if this were not the case then the resistant variety would be preferable in all situations). Tolerance, denoting resilience to viral load with minimal yield loss [30], is reflected in an *αβ* value for *TOL* that is equivalent to that of *SUSC* but with a smaller yield loss ratio – just 10% of that of *SUSC*. Hence, *TOL* experiences lower yield loss but also has a lower potential yield (*Y*_*h*_), preventing it from systematically outperforming *SUSC* in terms of yield when grown in mixtures.

In addition, to the main phytotypes we also consider the possibility of intercropping using a non-host crop which is a member of the host set for the polyphagous cassava whitefly [27, 13]. Hence, we consider a non-host ‘decoy crop’ (*DEC*) that does not produce cassava tubers nor acquire or transmit the virus. It is therefore characterized by *α* = *β* = *Y*_*h*_ = *Y*_*i*_ = 0.

#### Cassava Phytotype profiles for Cassava Mosaic Disease

The baseline values of *α* and *β* for CMB are estimated in Donnelly et al. [9] as *α* = 0.2688 day^−1^ and *β* = 132.21 day^−1^. The reference product *αβ* is 35.54 day^−2^, making it 3.54 for *RES*. We attribute the *Y*_*h*_ and *Y*_*i*_ values based on the literature of CMD, considering all CMB strains equally.

1. To create a *SUSC* profile, we consider a local susceptible variety in Côte d’Ivoire, Bonoua, whose average yield is approximately 21 tons/ha [21]. This variety is grown in Bonoua in the south, where the average CMD incidence is about 50% [20].
2. We assume most of the yield loss is associated with CMD so that the total yield is 21 = 0.5×*Y*_*h*_ +0.5×*Y*_*i*_.
3. Moreover, Samura et al. [28] estimate that full disease leads to a 40% yield loss, from which *Y*_*i*_ = 0.6 × *Y*_*h*_. Thus, *Y*_*h*_ *∼* 31 tons/ha and *Y*_*i*_ = 18.6 tons/ha.
4. Resistant varieties can have low yields, e.g., TME7 (22.85 tons/ha) after 12 months Bakayoko et al. [1], or higher. In accordance, we consider that the *RES* profile yield about 20% less than *SUSC, Y*_*h*_ = 25 tons/ha, and applying the same 40% yield loss results in *Y*_*i*_ = 15 tons/ha.
5. *TOL* has a reference yield of *Y*_*h*_ = 25 tons/ha, like *RES*, but experiences only an 10% less yield when infected, resulting in *Y*_*i*_ = 24 tons/ha.

We summarize the cassava phytotype profiles for CMD in Table2 A.

**Table 2:** Four cassava phytotypes for CMD and equivalent phytotypes for CBSD.

| A, CMD phytotypes | Phytotype name | Epidemiological parameters |  | Yield parameters |  |
| --- | --- | --- | --- | --- | --- |
| | | $\alpha$ | $\beta$ | $Y_h$ | $Y_i$ |
| Susceptible phytotype | *SUSC* <sup>(1)</sup> | 0.27 day<sup>-1</sup> | 132.21 day<sup>-1</sup> | 31 ton/ha | 18.6 ton/ha |
| Resistant phytotype | *RES* <sup>(2)</sup> | 0.09 day<sup>-1</sup> | 41.81 day<sup>-1</sup> | 25 ton/ha | 15 ton/ha |
| Tolerant phytotype | *TOL* <sup>(3)</sup> | 0.27 day<sup>-1</sup> | 132.21 day<sup>-1</sup> | 25 ton/ha | 24 ton/ha |
| Decoy phytotype | *DEC* | 0.0 day<sup>-1</sup> | 0.0 day<sup>-1</sup> | 0 ton/ha | 0 ton/ha |
| B, CBSD phytotypes | Phytotype name | $\alpha$ | $\beta$ | $Y_h$ | $Y_i$ |
| Susceptible phytotype | *SUSC* <sup>(4)</sup> | 68.88 day<sup>-1</sup> | 1.19 day<sup>-1</sup> | 31 ton/ha | 3.1 ton/ha |
| Resistant phytotype | *RES* <sup>(5)</sup> | 21.78 day<sup>-1</sup> | 0.38 day<sup>-1</sup> | 25 ton/ha | 2.1 ton/ha |
| Tolerant phytotype | *TOL* <sup>(6)</sup> | 68.88 day<sup>-1</sup> | 1.19 day<sup>-1</sup> | 25 ton/ha | 22.75 ton/ha |
| Decoy phytotype | *DEC* | 0.0 day<sup>-1</sup> | 0.0 day<sup>-1</sup> | 0 ton/ha | 0 ton/ha |

<sup>(1)</sup> CMB virological parameters were estimated in Donnelly et al. [9]. The reference yield when healthy ( $y_h$ ) is computed from Kouassi Kan et al. [21], Kouakou et al. [20] and the yield when infected ( $y_i$ ) is computed from Samura et al. [28] as 60% of $y_h$ .
<sup>(2)</sup> The product $\alpha\beta$ is determined as 10% of the reference *SUSC*'s. The yield when healthy is considered from Bakayoko et al. [1] and $y_i$ is computed by applying 40% of yield loss as in *SUSC*.
<sup>(3)</sup> The inoculation and acquisition rates are the same as for *SUSC*. The yield when uninfected is the same as for *RES*, and only 4% of yield is lost when infected, compared to 40% in others.
<sup>(4)</sup> Epidemiological parameters are estimated from Donnelly et al. [9] and yield loss is 90% for CBSD.
<sup>(5)</sup> The product $\alpha\beta$ is determined as 10% of the reference *SUSC*'s and yield loss remains 90%.
<sup>(6)</sup> The epidemiological parameters are considered the same as for *SUSC*. The yield when uninfected is the same as for *RES*, and only 9% of yield is lost when infected, compared to 90% in others.
– See more details in-text.

#### Cassava Phytotypes for Brown Streak Disease

We assume an extension to CBSD by assuming yield losses of 90% in susceptible and resistant varieties and 9% in tolerant varieties. Epidemiological parameters *α* and *β* are derived from CMD phytotypes by multiplying by the CMD-to-CBSD ratio in Donnelly et al. [9]. These multiplication factors are 256.25 for *α* and 9.08 10^−3^ for *β*, which leads to baseline values *α* = 68.88 day^−1^, *β* = 1.19 day^−1^ and the product *αβ* = 81.96 day^−2^. Note that this product is higher than in CMD, but conversely, here the insects recover with a rate *r* = 28.08 day^−1^ [9].

We also summarize the cassava phytotype profiles for CBSV in Table2.

#### Additional Parameters

Independently of the plant phytotypes, a few parameters need to be defined (Table 3).

**Table 3:** Parameter values. ^(*a*)^ is the median value from the posterior dispersal distribution in Ferris et al. [12]. ^(*b*)^ The vector death rate was estimated between 0.06 and 0.18 [18]. The growth rate of vectors was estimated at 0.2 based on laboratory experiments. However, analysis of natural population curves revealed a slope of 0.0118, suggesting that mortality contributes approximately 0.1882 (i.e. 0.2 − 0.0118) to the overall growth rate [36]. Bayesian parameter estimation and hypothesis testing have shown that the available laboratory data does not support whitefly recovery from CMD ^(*c*)^, and that the whitefly median duration of infectiousness with CBSD is approximately 1 hr ^(*d*)^ is the median value from the posterior recovery distribution in Donnelly and Gilligan [10], Donnelly et al. [9]. ^(*e*)^ Cassava takes an average of 10-12 months to mature before harvest but, in some cases, cassava can take up to 24 months to reach full maturity [3]. ^(*f*)^ is a representative choice based on literature ranges [12].

| Notation | Parameters | Value |
| --- | --- | --- |
| $\sigma$ | Insect dispersal | 0.45 days<sup>-1</sup> <sup>(a)</sup> |
| $\omega$ | Insect mortality | 0.19 days<sup>-1</sup> <sup>(b)</sup> |
| $r$ | Insect recovery from infectiousness | 0 days<sup>-1</sup> <sup>(c)</sup> (CMD)<br>28.08 days<sup>-1</sup> <sup>(d)</sup> (CBSD) |
| $T$ | Cropping duration | 360 days <sup>(e)</sup> |
| $\gamma$ | Incubation duration | 1/30 days<sup>-1</sup> <sup>(f)</sup> |

### 2.4 Insect Burden and Health Status

In our case study, we define insect burden through the vector abundance per plant. We define 3 different cases: (1) low insect burden (1 insect per plant), (2) medium insect burden (5 insects per plant), (3) high insect burden (10 insects per plant). We assume that the field is initially disease-free, and we consider an initial inoculum of 1 infected plant, while the vectors are all initially virus-free. Since it is challenging to decide which variety is initially infected, especially when the mixture is monocultural, we considered *θ* infected variety A plants and (1 − *θ*) infected variety B plants, which makes sense in a mean-field model. If one of the varieties is *INT*, then the infected plant is in the other variety.

### 2.5 App Implementation and Visualization

The mathematical framework is implemented as a user-friendly decision support tool (DST) in the form of a web application: CropMix (https://mixture-simulator.streamlit.app/). The DST follows a design framework structured around three core components: inputs, outputs, and internal parameters. The App implementation and a use case example are summarized in Box 2

#### Box 2: Crop Mix design – https://mixture-simulator.streamlit.app/

**Framework and implementation**

The app design revolves around the following key components:

1. Inputs: Include a disease to analyze, the insect burden and control inputs: cultivar mixture levels of interest and roguing rate.
2. Internal Parameters: Factors like infection rates and disease progression, which are crucial for modelling disease dynamics, and are set in the app’s settings.
3. Outputs: (i) Disease dynamics, infected insect dynamics, and resulting yields, and (ii) suggestions for the optimal mixture upon request, along with the associated yields and disease and insect dynamics.

**Figure 3:**
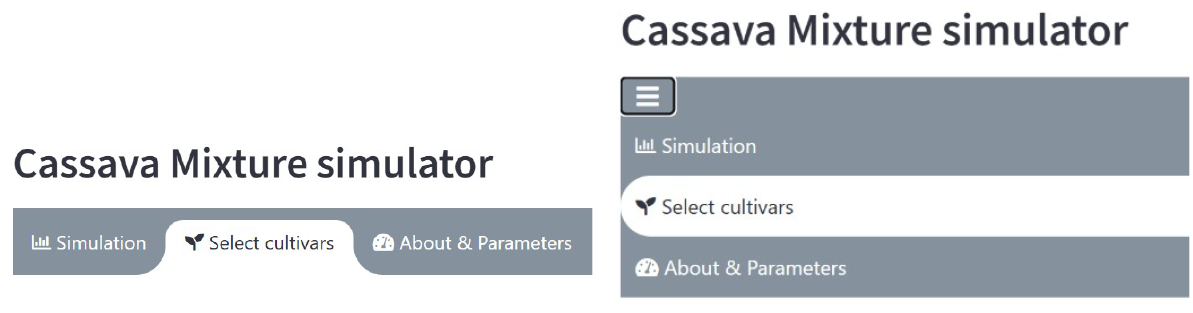
Desktop and mobile views of the simulation app tabs. CropMix is organized into three tabs: (1) a Simulation tab where the user can set the input and observe the outputs; (2) a cultivar selection tab (“Select Cultivars”) where the user selects two varieties for analysis, each represented by a plant phytotype; (3) a settings tab (“About and parameters”) that provides an overview of the model and allows the user to edit intrinsic parameters.

**Example Use Case**

To study a mixture for CBSV under moderate vector abundance, we selected two cultivars (‘Susceptible’ and ‘Resistant’) in the Select Cultivar Tab from the options: ‘Susceptible,’ ‘Resistant,’ ‘Tolerant’, and ‘Decoy’. We then returned to the Simulation Tab, where we selected ‘CBSD’ from the disease menu and set the vector abundance input to ‘medium’ in the plant-wise insect burden menu. By default, the app displayed yield, disease dynamics, and infected insect dynamics over time for a monoculture of ‘Susceptible.’ We adjusted the mixture slider to observe outputs for different mixtures and set a roguing frequency to examine the combined effects of cultivar mixture and roguing. After selecting a specific roguing frequency (every 30 days), we clicked the “Find the Optimal checkbox”, allowing the app to discover the optimal mixture, which resulted in a 564:436 ratio of Susceptible and Resistant. This means it is optimal to grow 5,640 susceptible plants and 4,360 resistant plants in a hectare with a density of 10,000 plants per hectare.

## 3 Results

The results displayed here are based upon the web application output for particular scenarios (section 2.5). They explore strategies that protect susceptible cassava *SUSC* against cassava mosaic disease (CMD) and cassava brown streak disease (CBSD), or that mitigate losses from these diseases, under varying levels of whitefly pressure. In summary, our analyses show that no mixture is able to fully protect *SUSC* from CMD. For CBSD, however, while resistant cassava *RES* can often protect *SUSC*, its effectiveness diminishes under high insect pressure. The decoy crop *DEC* can significantly improve *SUSC* yield by providing plants for virus-bearing whiteflies where transmission of the virus does not occur especially under moderate whitefly pressure (less *DEC* is required for protection so the yield penalty is lower). However, under certain conditions, our analyses indicate that regular roguing can actually eliminate the need for *DEC*. In addition, tolerant varieties make sense only as mono-cultures - in mixtures they will act as sources of virus for the susceptible component of the mixture - and in general the optimality of tolerant mono-cultures depend upon their particular attributes. Overall, these aspects illustrate that combining control strategies like roguing with crop mixtures can be an effective means of disease management with mono-cultures of tolerant varieties an important option when such management practices are not sufficiently effective. These results are explored in more detail below.

### 3.1 Cassava Mosaic Disease

#### No mixture can protect susceptible plants against CMD

No mixture can protect susceptible plants against CMD, making monocultures the optimal choice in every scenario. Under low insect pressure, disease incidence quickly rises to 100% (Fig 4), making the *TOL* monoculture the optimal choice, regardless of the presence of other varieties. Consequently, this monoculture remains optimal even as insect pressure increases (medium, high). When roguing is introduced as an additional control method, the *SUSC* monoculture may become optimal, depending on the frequency of roguing, as disease incidence rise to lower levels than 100% (Fig 4). Since the optimal configurations are always monocultures, there is no need to evaluate *DEC*, which provides no yield. Table 4 summarizes the optimal monocultures across various scenarios.

**Table 4:** No mixtures can protect susceptible cassava from CMB: Optimal monocultures for diverse setups. When no roguing is performed or when roguing is less frequent than every 3 weeks, the tolerant monoculture yields the best, no matter the insect burden. Conversely, with more frequent roguing, the susceptible monoculture yields the best. The resulting yields vary little with the insect burden. ^(*a*)^ The yields are calculated *ρ* = 1/30 and are slightly lower for *ρ* < 1/30 while the order of best yielding monocultures remains unchanged. ^(*b*)^ The yields are calculated for *ρ* = 1 14 and are slightly higher for *ρ* > 1 14 while the order of best yielding monocultures remains unchanged.

| Roguing Frequency | Insect burden |  |  |
| --- | --- | --- | --- |
|  | Low | Medium | High |
| None | *TOL* (24 ton/ha) |  |  |
|  | *SUSC* (18.6 ton/ha) |  |  |
|  | *RES* (15 ton/ha) |  |  |
| 1 month or more<br>between roguing rounds (roguing rate $\rho \leq 1/30$) <sup>(a)</sup> | *TOL* (24.3 ton/ha) | | |
|  | *SUSC* (22.6 ton/ha) |  |  |
|  | *RES* (18.2 ton/ha) |  |  |
| 2 or less weeks<br>between roguing rounds (roguing rate $\rho \geq 1/14$) <sup>(b)</sup> | *SUSC* (24.8 ton/ha) | | |
|  | *TOL* (24.5 ton/ha) |  |  |
|  | *RES* (20 ton/ha) |  |  |

**Figure 4:**
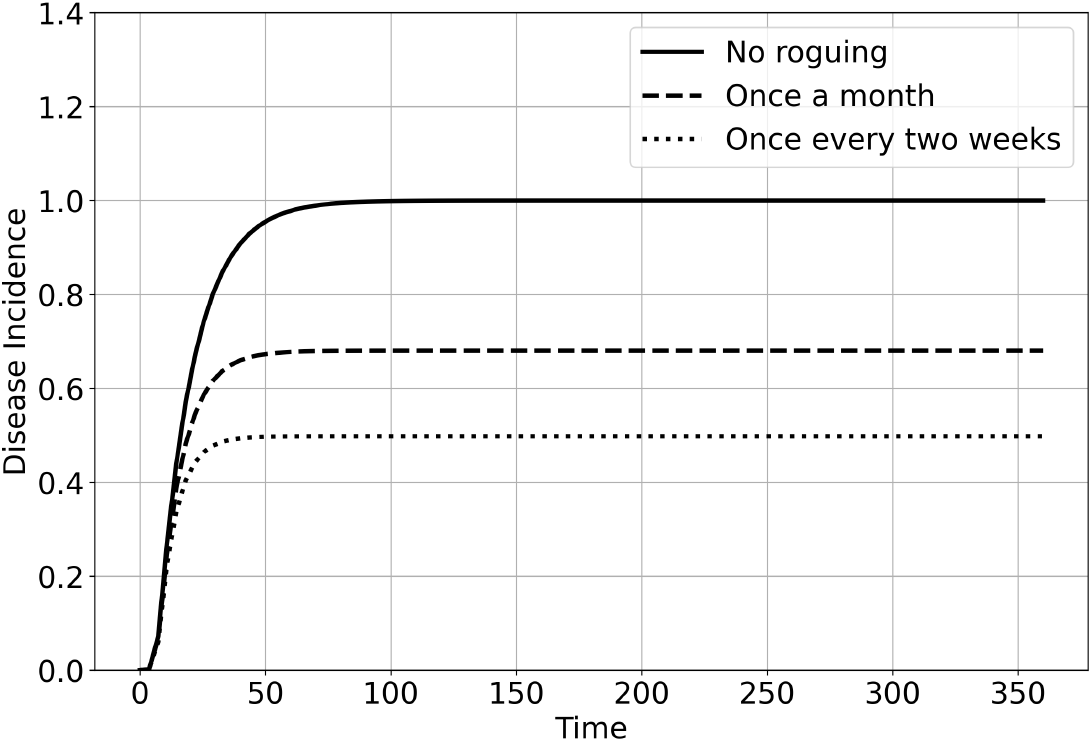
CMD incidence dynamics under low insect pressure for three different roguing frequencies: no roguing, once a month, and once every two weeks. In the absence of roguing, incidence rapidly rises to 100%, whereas the saturation level decreases as roguing becomes more frequent.

### 3.2. Cassava Brown Streak Disease

Table 5 and Fig 10 summarize the mixtures that optimize cassava yields in a field infected by CBSD for diverse setups. In subsequent subsections, we analyze the outcomes for actual cassava mixtures first, then for the decoying protection *DEC* might offer to *SUSC*.

**Table 5:** Optimal mixtures against CBSD for diverse setups. The table presents: (i) the reference yield of *SUSC* grown alone, (ii) the optimal mixture of *SUSC* with RES and its yield, (iii) the optimal mixture of *SUSC* with *TOL* and its yield, and (iv) the optimal mixture of *SUSC* with *DEC* and its yield. The latter three optimal mixtures may be monocultures. When the optimal mixture consists solely of *SUSC*, as in the case of mixing with *DEC*, an additional row is unnecessary. ^(*a*)^ The yields are calculated for *ρ* = 1/30 and are lower for *ρ* < 1/30. The optimal mixture proportions may vary with reduced values of the roguing rate *ρ* until they stabilize to the same profile as when there is no roguing.^(*b*)^ The yields are calculated for *ρ* = 1/14 and are higher for *ρ* > 1/14 than for *ρ* = 1/14. For higher values of the roguing rate *ρ*, the best-yielding order remains unchanged: the *SUSC* monoculture remains optimal under low and medium insect burdens, while the *SUSC*–*RES* mixture remains optimal under high insect burden. In the latter case, the proportion of *SUSC* in the mixture increases with the roguing rate, reaching 100% as *ρ* approaches 1 (roguing every day), as illustrated in Supplementary Material S5. More generally, increasing the frequency of roguing increases the share of *SUSC* in mixtures.

| Roguing Frequency | Insect burden |  |  |
| --- | --- | --- | --- |
|  | Low | Medium | High |
| None | *SUSC*<br>(30.3 ton/ha) | 10.6% *SUSC* - 89.4% *RES*<br>(25.1 ton/ha) | *RES*<br>(22.5 ton/ha) |
|  | *RES*<br>(25 ton/ha) | *TOL*<br>(22.7 ton/ha) | *TOL*<br>(22.7 ton/ha) |
|  | *TOL*<br>(25 ton/ha) | 38.5% *SUSC*- 61.5% *DEC*<br>(9.5 ton/ha) | 17.1% *SUSC*- 82.9% *DEC*<br>(4.1 ton/ha) |
|  |  | *SUSC*<br>(3.1 ton/ha) | *SUSC*<br>(3.1 ton/ha) |
| 1 month or more<br>between roguing rounds<br>(roguing rate $\rho \leq 1/30$) <sup>(a)</sup> | *SUSC*<br>(31 ton/ha) | 90.2% *SUSC* - 09.8% *RES*<br>(29.8 ton/ha) | 30.3% *SUSC* - 69.7% *RES*<br>(26.5 ton/ha) |
|  | *RES*<br>(25 ton/ha) | *SUSC*<br>(29.5 ton/ha) | *TOL*<br>(23.8 ton/ha) |
|  | *TOL*<br>(24.99 ton/ha) | *TOL*<br>(24.9 ton/ha) | *SUSC*<br>(16.8 ton/ha) |
| 2 or less weeks<br>between roguing rounds<br>(roguing rate $\rho \geq 1/14$) <sup>(b)</sup> | *SUSC*<br>(31 ton/ha) | *SUSC*<br>(31 ton/ha) | 75.9% *SUSC* - 24.1% *RES*<br>(29.15 ton/ha) |
|  | *RES*<br>(25 ton/ha) | *RES*<br>(25 ton/ha) | 94.3% *SUSC* - 5.7% *DEC*<br>(26.14 ton/ha) |
|  | *TOL*<br>(25 ton/ha) | *TOL*<br>(25 ton/ha) | *SUSC*<br>(26.1 ton/ha) |
|  |  |  | *TOL*<br>(24.6 ton/ha) |

#### Cassava mixtures

At low levels of insect pressure, disease incidence is so low that yield loss for all varieties is marginal. As a result, *SUSC* dominates any mixtures it is involved in. In such cases, mixtures are unnecessary to protect *SUSC*. Conversely, under moderate to high insect pressure, *TOL* never emerges as being beneficial in mixes with *SUSC* because the yield loss due to CBSD is generally so high that *TOL* dominates any potential mixture.

Under moderate insect pressure, the optimal configuration is the *SUSC-RES* mixture (Fig 5), which yields over 700% more than when *SUSC* is not protected by *RES*. If a higher proportion of *SUSC* is required in the mixture, due for example to market demand or food quality considerations, our model results suggest that regular roguing could help. For instance, roguing once a month increases the proportion of *SUSC* in the mixture to 90.2% (Fig 6), achieving a yield of 29.8 tons/ha, which surpasses the yield without roguing (25.1 tons/ha). Notably, the monoculture of *SUSC* under monthly roguing is also highly productive (29.5 tons/ha), underscoring that roguing benefits *SUSC*.

**Figure 5:**
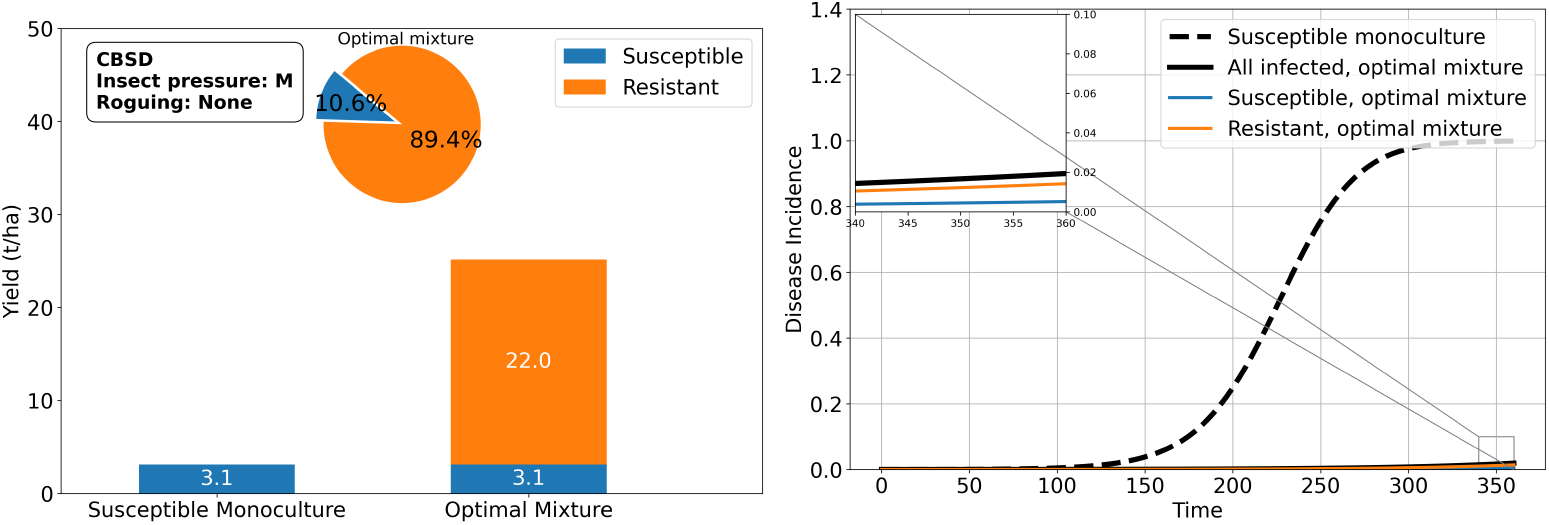
A susceptible-resistant mixture (10.6% susceptible vs. 89.4% resistant) is yield-wise optimal under moderate whitefly pressure and brown streak disease. This mixture produces a yield of 22.0 tons/ha for the resistant variety and 3.1 tons/ha for the susceptible variety. Notably, the susceptible cassava yield in the mixture matches that of a susceptible monoculture, despite the smaller proportion of susceptible cassava (10.6%) in the mixture. Disease incidence is significantly reduced in the mixture.

**Figure 6:**
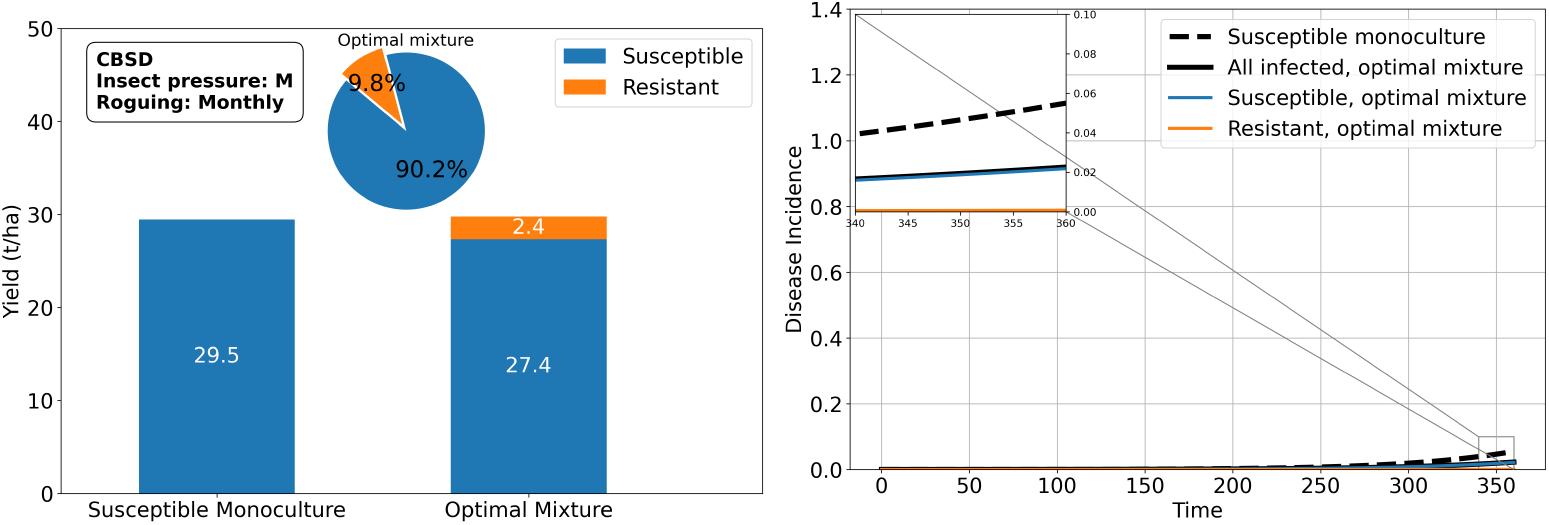
Roguing increases the proportion of the susceptible cassava variety in the optimal mixture with the resistant variety under moderate insect pressure and brown streak disease. When roguing is performed once a month, the optimal mixture contains 9.8% resistant plants, compared to 82.9% in the absence of roguing. Overall, the total yield with roguing in the optimal mixture (29.8 tons/ha) is slightly higher than in a susceptible monoculture (29/5 tons/ha), and both are significantly better than scenarios without roguing, due to the reduced disease incidence.

Under high insect pressure, our results indicate the best configuration is a monoculture of *RES*, as neither *RES* nor *TOL* effectively protects *SUSC* in mixtures. However, if *SUSC* is highly desired, decoying strategies (as we explore in the next result) or roguing can help reintroduce *SUSC*. With monthly roguing, the optimal mixture comprises 30.3% *SUSC* (Fig 7), yielding more than monocultures of *SUSC, TOL*, or *RES* alone. More generally, roguing promotes the share of *SUSC* in the mixtures.

**Figure 7:**
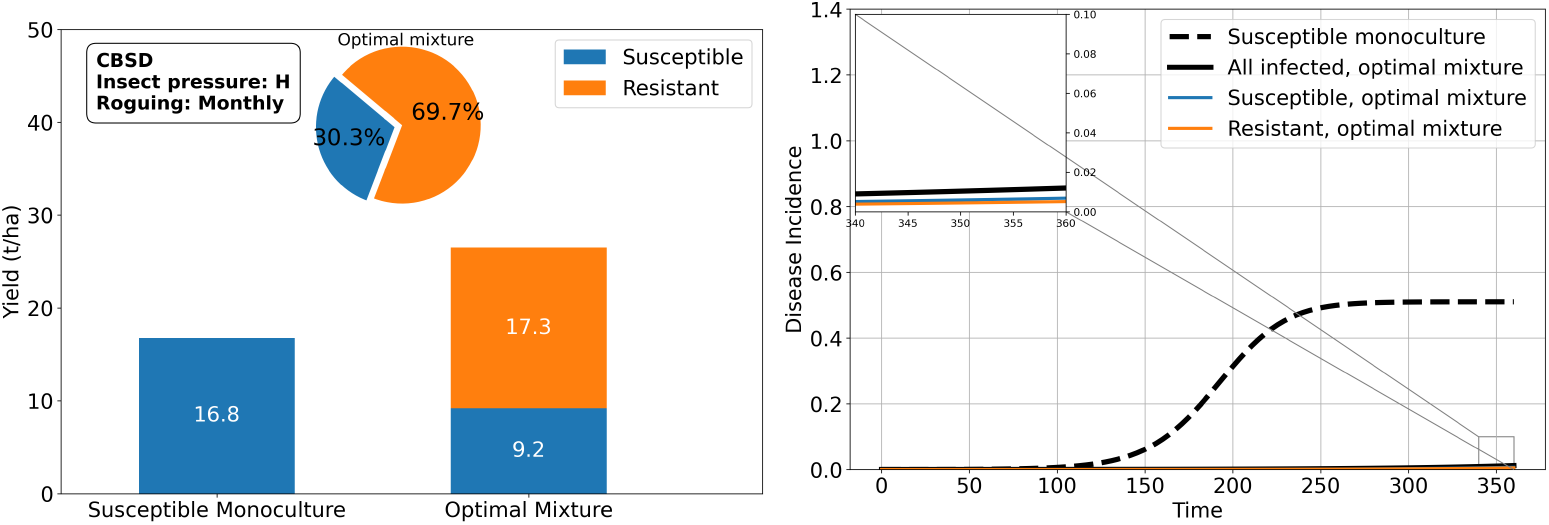
Roguing reintroduces the susceptible cassava variety in the optimal setup under high insect pressure and brown streak disease: a proportion of 69.7% of resistant plants is optimal in a mixture with susceptible when roguing is performed once a month, while the resistant monoculture was optimal in the absence of roguing.

**Figure 8:**
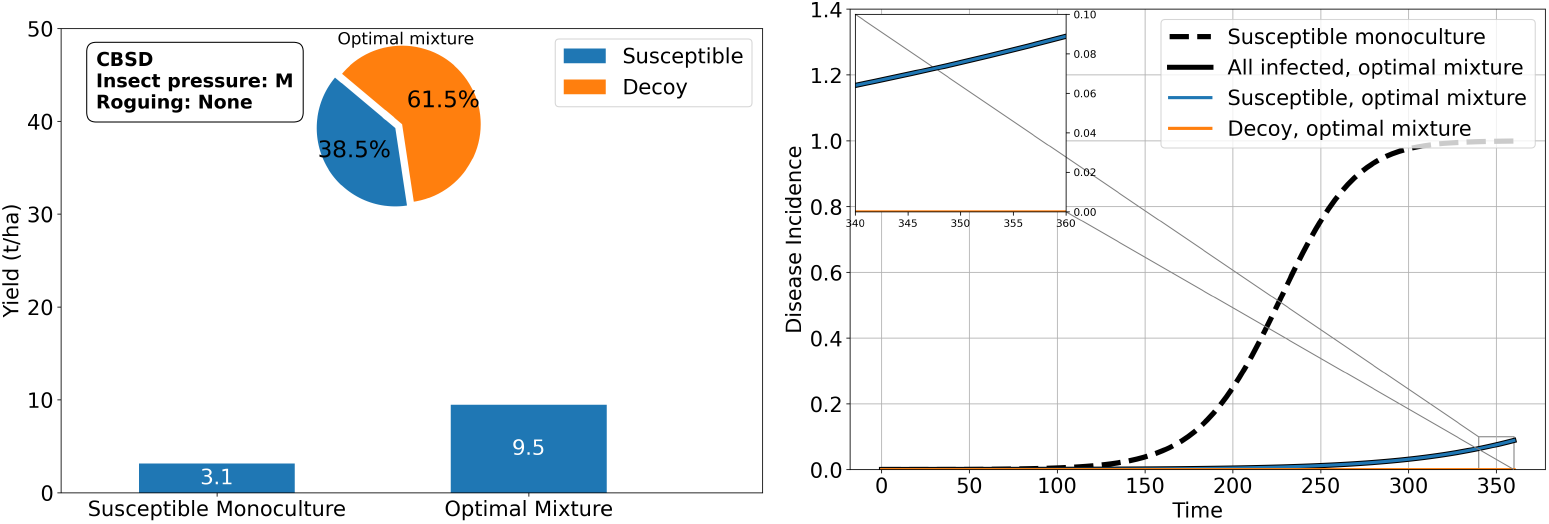
Decoy plants can protect cassava from cassava brown streak ipomovirus under moderate whitefly pressure. An optimal proportion of 61.5% decoy plants attracts enough whiteflies to significantly lower the disease incidence. This insures 9.5 tons/ha of cassava yield associated with the 38.5% of susceptible cassava in the mixture, while a monoculture of susceptible cassava achieves 3.1 tons/ha.

These results highlight that *RES* consistently outperforms *TOL* in protecting *SUSC* in mixtures when *SUSC* is desired. However, under extremely high insect pressure, *RES* may fail to safeguard *SUSC*, which can be problematic if *SUSC* is in high market demand. Fortunately, decoy crops can be used to reintroduce *SUSC* and meet market needs, as further analyzed in the next subsection.

#### Insect decoying

As shown in the previous results, *RES* is often sufficient to protect *SUSC* from CBSD and achieve high yields. However, under high insect pressure, *RES* may fail to provide adequate protection. In such cases, growing *DEC*, which is opaque to the virus but attracts whiteflies similarly to cassava, can help support the cultivation of *SUSC*. Our results indicate that mixing *DEC* with *SUSC* can yield up to 4.1 tons per hectare when 82.9% of the field is planted with *DEC*. This represents a 32% improvement compared withgrowing *SUSC* alone (Fig 9). Remarkably, even with *SUSC* occupying only 17% of the field, the yield is significantly higher than planting *SUSC* exclusively, provided the *SUSC* crops are protected by *DEC*.

**Figure 9:**
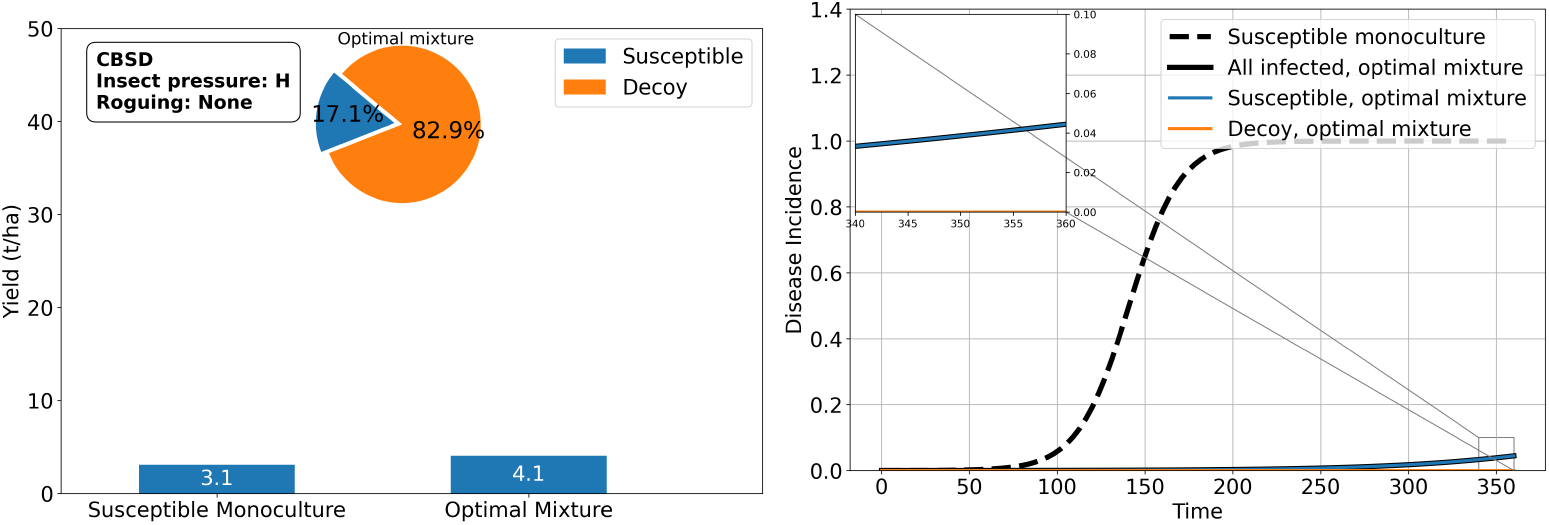
Decoy plants can protect cassava from cassava brown streak ipomovirus under high whitefly pressure. An optimal proportion of 82.9% decoy plants attracts enough whiteflies to significantly lower the disease incidence. This insures 4.1 tons/ha of cassava yield associated with the 17.1% of susceptible cassava in the mixture, while a monoculture of susceptible cassava achieves 3.1 tons/ha.

Under moderate whitefly pressure, the protective effect of *DEC* is even greater, enabling up to a 200% improvement in *SUSC* yield with a smaller proportion of *DEC* required (61.5%, see Fig 8). Additionally, *DEC* enhances the productivity of *SUSC* compared with a mixture with *RES* (9.5 tons/ha vs. 3.1tons/ha), despite the fact that a *RES* monoculture remains optimal in terms of total yield (25.1 tons/ha vs. 9.5 tons/ha).

With roguing, however, the need for *DEC* vanishes. Therefore, mixing *DEC* with cassava varieties can be just considered an additional control strategy, complementary to varietal mixtures.

#### Summary

In Fig 10 we summarise the yields of optimal setups across each kind of mixture, which is a graphical representation of Tables 4 and 5.

**Figure 10:**
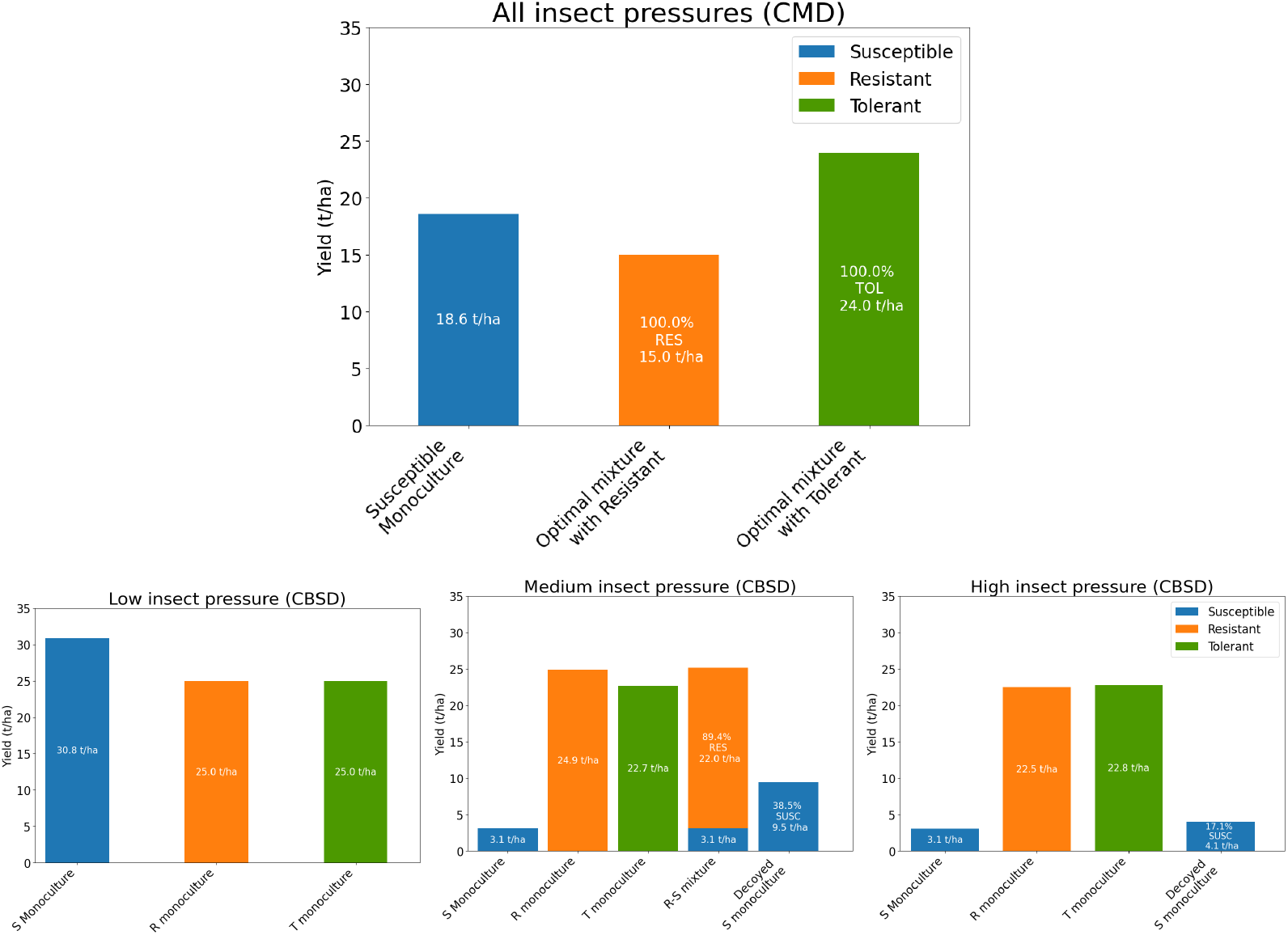
Summary - Cassava yields in diverse growing scenarios without roguing in the presence of CMD (top) and CBSD (bottom). Regardless of insect pressure, the incidence of CMD quickly rises to 100%, and the tolerant monoculture yields the best as it responds better to disease-induced yield loss. Susceptible varieties do not need resistance shielding from CBSD under low whitefly pressure, but as insect pressure increases, resistant varieties become more dominant. As a result, a mixture of susceptible and resistant varieties is optimal under moderate whitefly pressure, while a monoculture of resistant varieties is optimal under high whitefly pressure. In both cases, shielding susceptible cassava crops with decoying plants that attract whiteflies similarly is more effective than growing susceptible cassava alone, despite the significantly reduced proportion of cassava grown.

## 4 Discussion

In this work, we have presented optimal crop mixture results for epidemic scenarios relating to cassava mosaic disease (CMD) and cassava brown streak disease (CBSD) infection. The results are based on output for these scenarios from the novel web application CropMix which is introduced in this paper. The web application is designed to receive input relating to attributes of a number of cultivars and to calculate varietal mixture levels that optimise yield in the face of crop disease. Taking the motivating example of whitefly-borne cassava viruses, we use the approach to investigate the potential for virus-resistant cassava to protect virus-susceptible cassava. Our results indicate that broadly that the protective effect of resistant cassava is feasible for CBSD but not for CMD. We also consider varieties that are tolerant to viral disease as well as intercropping with alternative crops that are within the host range of the polyphagous cassava whitefly. Overall, our results indicated that none of the CMD-resistant cassava phytotypes that were tested could protect CMD-susceptible cassava. Instead, deployment of CMD-tolerant monocultures, and to a lesser extent CMD-resistant monocultures, are more likely to produce reasonable crop yields. This is due to the very high transmissibility of CMB; varietal mixing and intercropping were unable to prevent the rapid rise in disease incidence. Our results indicated that CBSD-resistant cassava, however, could protect CBSD-susceptible cassava, with the results holding across a wide range of scenarios. When insect pressure is very high, however, the high transmission that results means that cropping with CBSD-tolerant monocultures and to a lesser extent CBSD-resistant monocultures, are more likely to produce reasonable crop yields (as was the case more generally with CMD). It is also important to consider the degree to which non-host crops can protect disease-susceptible varieties. For insect-borne plant pathogens, this works by providing decoy plants for virus-bearing whiteflies in which transmission of the virus does not occur. The strategy requires decoy plants to have a similar insect vector affinity – and for this reason, our investigations are relevant for crop plants within the host set of the polyphagous cassava whitefly. On the whole, we found that this type of intercropping performed well in the situations in which susceptible-resistant mixtures performed well. Thus the susceptible portion of planted fields produced a higher yield than without an alternative crop: this is true even without considering the alternative yield from the non-host crop. These findings align with prior research demonstrating that intercropping enhances resilience by disrupting pathogen transmission and reducing disease severity [27, 13].

### Additional considerations

It is important to contextualise the results presented in the paper. Policy-makers in particular should bear the following points in mind. Firstly, tolerant varieties make sense only as mono-cultures: in mixtures, they are likely to act as vigorous sources of virus for susceptible varieties. Furthermore, regions that propagate tolerant mono-cultures may prove significant sources of virus for surrounding areas. As such, while tolerant varieties are highly valuable it is important to first assess whether or not alternative strategies like varietal mixing and intercropping offer viable disease management options. Our results pertaining to intercropping with alternative non-host crops assume identical insect vector affinity on the non-host crop as compared with the main host crop. Additional feasibility studies are needed to quantify this affinity for candidate crops.

As already discussed we found that mixtures and inter-cropping were ineffective for managing CMD. So what can be done to combat this important disease? Roguing has sometimes proven effective in reducing inoculum sources for CMD leading to decreased disease incidence. All of this suggests that additional management practices like roguing can complement varietal mixtures by further curbing disease spread, emphasizing the value of integrated strategies. Similar findings are often noted in other pathosystems, where integrating cultural practices enhances yields [22]. Studies have shown that timely and thorough roguing significantly delays disease spread and it was advised that regular roguing, performed at least every week, could lower disease incidence and sustain yields in cassava fields [33]. However, roguing success depends on early detection, proper timing, and the thoroughness of the process, as delayed or incomplete roguing may leave sufficient viral inoculum to perpetuate outbreaks. As a caution, roguing is ineffective and counterproductive in areas with high virus pressure, as it reduces plant populations to unacceptable levels, which our model does not capture as systematic replanting is assumed.

To improve the precision and applicability of the model, integrating real-world plant varieties, rather than relying solely on generalized phytotypes, is an important next step. This would involve a more detailed parameterization process, including gathering variety-specific data on key factors including yields, virus acquisition and inoculation rates, and incubation periods. Laboratory-based experiments would be crucial for determining these transmission parameters for each variety [11, 10]. Conducting comprehensive field trials to quantify yield losses under varying levels of disease incidence across different varieties and environmental conditions is vital. Such data would allow for updates to the model’s internal parameters, ensuring that it remains aligned with field systems without requiring significant structural changes.

## 5 Conclusion

Mixtures and inter-cropping were ineffective for managing CMD but can significantly mitigate the impact of CBSD. In particular, CBSD yield is preserved by resistant varieties shielding susceptible varieties under varying levels of insect pressure. In situations where the degree of virus transmission is too high and resistant-susceptible mixtures are not profitable, tolerant mono-cultures may be optimal but only if they have the right attributes in terms of satisfactory yield returns. Nevertheless, tolerant mono-cultures do not help the epidemiological situation regionally and so due consideration should be given to intercropping with non-host crops for which the insect vector has an affinity.

## Notes

### Competing Interest Statement

The authors have declared no competing interest.

